# Voluntary oral fentanyl intake produces dose- and sex-dependent physical dependence in mice without overt affective disturbances

**DOI:** 10.64898/2026.07.06.736848

**Authors:** Marie-Charlotte Allichon, Samuel F Boehm, Nilah D Jordan, Lars H Nelson, Max E Joffe

**Affiliations:** Center for Neuroscience, University of Pittsburgh, Pittsburgh, PA 15219, USA; Translational Neuroscience Program, Department of Psychiatry, University of Pittsburgh, Pittsburgh, PA 15219, USA

**Keywords:** oral self-administration, opioid use disorder (OUD), sex differences, rodent model, polysubstance use

## Abstract

The ongoing opioid epidemic underscores the need for scalable and translational preclinical models of voluntary opioid intake and dependence. We therefore sought to establish and validate a voluntary two-bottle choice drinking-in-the-dark (DID) model of oral opioid intake in mice and to determine relationships between experimental parameters and behaviors during and after withdrawal. Male and female C57BL/6J mice were given daily access to two bottles during the dark phase for 24 drinking sessions over 5 weeks. Control mice received two bottles containing water. Experimental mice received one water bottle and one bottle containing oxycodone (0.1-1 mg/mL) or fentanyl (10-100 µg/mL) under varying session durations and concentrations. On the final day, physical dependence was assessed using naloxone-precipitated withdrawal and then a behavioral battery to assess negative affect was performed in the following week. Mice voluntarily consumed both oxycodone and fentanyl without taste adulteration and maintained drug preference across most concentrations. Oxycodone intake produced minimal withdrawal symptoms. In contrast, fentanyl intake resulted in naloxone-precipitated withdrawal that was modulated by session duration and concentration. Four-hour sessions produced stronger withdrawal than two-hour sessions at equivalent concentrations. Escalating high-concentration fentanyl exposure revealed emerging sex differences, with females exhibiting greater intake and withdrawal at higher concentrations. Affective behavioral assays following withdrawal revealed minimal persistent alterations in any cohort. These findings establish key parameters for a scalable voluntary fentanyl model that produces dose- and session-dependent physical dependence in male and female mice. This paradigm provides a cost-effective and straightforward platform for future investigations of opioid use and dependence.

## Introduction

Opioid use disorder (OUD) represents a major public health crisis in the United States, affecting millions of individuals and driving substantial morbidity and mortality (CDC, 2025). Synthetic opioids such as fentanyl, a highly potent mu-opioid receptor agonist, are now among the most commonly misused substances. The rapid pharmacokinetic profile of fentanyl and related compounds promotes the development of compulsive use and increases the risk of dependence, as well as producing rapid respiratory depression and overdose risk (Comer and Cahill, 2019; Hill et al., 2020). In parallel, semi-synthetic opioids such as oxycodone remain widely prescribed analgesics while carrying significant risks of misuse and dependence (Cicero et al., 2007; Ordóñez Gallego et al., 2007; Drug Enforcement Administration, 2025). Understanding the neurobiological and behavioral mechanism that underlies susceptibility to develop OUD is therefore of critical importance. These ongoing public health challenges underscore the need for translational preclinical models that capture voluntary opioid intake and individual differences in the development of dependence.

Rodent models of opioid self-administration have traditionally relied on intravenous administration, which provides unparalleled control but requires catheterization and extensive training (Spealman and Goldberg, 1978). More recently, alternative routes including vapor (Marchette et al., 2023; Moussawi et al., 2020) and oral administration (Downs et al., 2024; Monroe and Radke, 2021) have been developed to increase scalability and accessibility. Similar self-administration approaches have been applied to oxycodone, demonstrating voluntary intake, escalation and the development in dependence in rodents (Enga et al., 2016; Iyer et al., 2022; Mavrikaki et al., 2017; Reeves et al., 2021; Slivicki et al., 2023; Zanni et al., 2021; Zhang et al., 2014). Notably, many of these studies were conducted exclusively in males, limiting our understanding of how sex can shape voluntary opioid intake, preference and dependence.

Oral self-administration paradigms are advantageous because there is no surgical burden and voluntary intake can be assessed over extended periods (Moore et al., 2007; Rhodes et al., 2005). Despite these strengths, previous studies using oxycodone without taste adulterants or forced exposure have not reliably found robust physical dependence (Iyer et al., 2022; Slivicki et al., 2023). More recent studies using fentanyl have been successful in eliciting dependence. Downs et al, 2024 (Downs et al., 2024) showed that oral fentanyl consumption produced naloxone-precipitated withdrawal; however, in these studies mice only had access to a single bottle with fluids, precluding assessments of voluntary intake and preference. Incorporating a two-bottle choice configuration within a time-restricted schedule enhances interpretability by distinguishing drug-directed preference from homeostatic fluid consumption (Sanchis-Segura and Spanagel, 2006).

Physical dependence following opioid exposure can be assessed using naloxone-precipitated withdrawal, which reliably elicits somatic signs such as escape jumps, paw tremors, and wet-dog shakes (Iwamoto et al., 1973; Lewter et al., 2022; Lichtman et al., 2001). While withdrawal following parenteral opioid administration is well-characterized, the parameters required for voluntary fentanyl intake to produce measurable withdrawal are not well-defined. A Drinking in the Dark (DiD) schedule, originally developed for binge-like ethanol consumption, restricts access to fluids during a set time in the dark phase (Rhodes et al., 2005; Thiele et al., 2014; Thiele and Navarro, 2014). The time-limited nature of DiD facilitates high levels of intake and facilitates investigations of behaviors that occur at discrete times, like withdrawal. Based on this, we aimed to determine ideal experimental parameters to elicit voluntary opioid intake, withdrawal severity, and affective behaviors in abstinence in both male and female mice

In the present study we implement a voluntary two-bottle DiD model of opioid intake in male and female C57BL/6J mice. We systematically manipulated session duration and concentration escalation to determine whether these parameters govern naloxone-precipitated withdrawal severity and related affective outcomes. Results are consistent with our hypotheses that (1) mice voluntarily consume dilute oxycodone and fentanyl without adulterants, (2) longer access and higher concentrations produce stronger withdrawal phenotypes, and (3) sex differences emerge under conditions of escalating fentanyl exposure. While we observed consistent intake over time and acute withdrawal behaviors, we did not observe clear or consistent affective behaviors during abstinence. Together, these findings provide a scalable and translationally relevant behavioral platform for studying voluntary opioid intake and physical dependence.

## Methods

### Animals

Adult (>10-week) C57BL/6J mice (males n = 42 and females n = 42) were used across four experimental cohorts. Determination of biological sex was limited to an assessment of external genitalia. Mice were bred at Charles River Laboratories, shipped to the University of Pittsburgh at 6-7-weeks, and acclimated to our facility for at least 2 weeks prior to investigation. Animals were pair-housed under a reverse 12/12 hour (lights off at 7:00am) with food and water always available ad libitum. All procedures were performed during the dark (active) phase. Body weight was measured after each drinking session. At study end animals were euthanized according to approved IACUC protocol.

### Drinking-in-the-Dark (DiD) Setup and Experimental Design

The DiD paradigm was adapted from published protocols (Downs et al., 2024; Iyer et al., 2022). At the start of each session, mice were transferred from the home cage to an experimental cage fitted with a perforated, transparent divider (Animal Care Systems), preserving social contact while allowing individual measurement (Peretz-Rivlin et al., 2025). Each mouse had access to two identical plastic bottles mounted side-by-side in housing adapted from open-source devices described previously (Godynyuk et al., 2019; Haggerty et al., 2025) (Supplemental Figure 3). Bottle positions were alternated across sessions to control for side bias. After the session mice were returned to the home cage. Body weight was recorded before each session.

### General experimental timeline

All cohorts underwent five weeks (Monday-Friday) of DiD sessions, one session per day, followed by a behavioral battery (see Figure 1). Monday of the first week served as a habituation session with two water bottles for all the animals. For each of the following 24 sessions mice were assigned within each cohort to one of two conditions: water/water (both bottles water) or drug/water (one bottle dilute opioid, one bottle water). Cohort-specific details follow.

**Figure 1.**
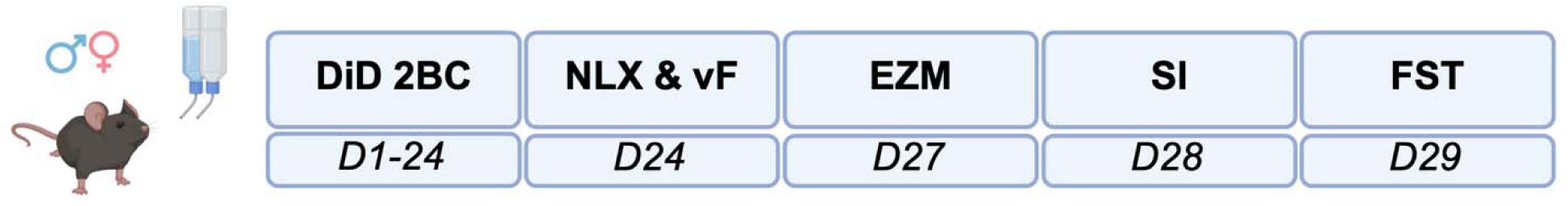
Standard experimental timeline used across all cohorts. Mice underwent 24 days of drinking in the dark (DiD) voluntary drug intake with a two-bottle choice (2BC) paradigm, followed on the final day of drinking by naloxone-precipitated withdrawal (NLX) assessment and von Frey (vF) testing. Affective-like behaviors were then assessed across the following three days. The Elevated Zero Maze (EZM) was conducted first, followed by the Social Interaction (SI) test and the Forced Swim Test (FST), with only one behavioral assay performed per day.

### Cohort specifics

Cohort 1: **Oxycodone 2 h/day.** n = 16 (8 males, 8 females). Session duration 4 h per day. Drug group received oxycodone HCl (Sigma, PHR8998) concentrations in water: 0.1 mg/mL on day 2, 0.3 mg/mL on day 3, 0.5 mg/mL on day 5, and 1 mg/mL from day 15 onward. Group composition: water group males n = 4 and females n = 4; oxycodone group males n = 4 and females n = 4.

Cohort 2: **Cohort 2: Fentanyl 2 h/day.** n = 20 (10 males, 10 females). Session duration 2 h per day. Drug group received fentanyl citrate (Sigma, PHR8977) concentrations in water: two days at 10 µg/mL, two days at 20 µg/mL, then 30 µg/mL for remaining sessions. Group composition: water group males n = 4 and females n = 6; fentanyl group males n = 6 and females n = 4.

Cohort 3: **Cohort 3: Fentanyl 4 h/day**. n = 24 (12 males, 12 females). Session duration 4 h per day. Same fentanyl concentration schedule as Cohort 2. Group composition: water group males n = 6 and females n = 6; fentanyl group males n = 6 and females n = 6.

Cohort 4: **Cohort 4: Fentanyl 4 h/day high-concentration.** n = 24 (12 males, 12 females). Session duration 4 h per day. Drug group received fentanyl citrate starting at 10 µg/mL and increasing by 10 µg/mL every two sessions up to 100 µg/mL, then maintained at 100 µg/mL for the remainder of the DiD. Group composition: water group males n = 6 and females n = 6; fentanyl group males n = 6 and females n = 6.

### Solution preparation and bottle handling

Opioid stock solution was prepared fresh at the beginning of the experiment and diluted to the appropriate concentration in drinking water. Bottles were checked for leakage; side positions were alternated for each session. Drug intake was calculated from bottle weights/volumes and reported as mg/kg for oxycodone and µg/kg for fentanyl using each animal’s body weight measured before the session.

### Naloxone-precipitated withdrawal

On day 24, each mouse received an injection of naloxone. Naloxone is a nonselective opioid receptor antagonist that elicits symptoms of withdrawal in opioid-dependent rodents (Downs et al., 2024; Iwamoto et al., 1973; Moussawi et al., 2020). First, animals received an intraperitoneal (i.p.) saline injection and were observed in a Plexiglass open field box (30 x 30 x 30 cm) for 20 minutes to obtain baseline behavior. Mice then completed their DiD session according to cohort timing and treatment. Immediately after the session mice were injected with naloxone HCl (2 mg/kg, i.p.; Hello Bio) and monitored in the same open field box for 20 minutes; sessions were recorded for offline scoring. Three stereotyped somatic withdrawal behaviors were scored from the 20-minute video: escape jumps, wet-dog shakes, and paw tremors.

### Von Frey mechanical sensitivity testing

Immediately following the withdrawal session mice were moved to a metal testing table for Von Frey evaluation following the SUDO simplified up-down method (Bonin et al., 2014). Animals were placed under an inverted cup and allowed 10 minutes to habituate. One hind paw was tested with calibrated monofilaments (BioSeb) held in place for 3 seconds per stimulus; a positive response (purposeful withdrawal, vigorous shaking, or licking) led to a weaker filament on the next trial, whereas no response led to a stronger filament. Five trials were conducted per animal with 5 minutes between trials. Paw withdrawal threshold (PWT) in grams was calculated using the SUDO-derived equation and the filament set constant, following this equation, where F is the log value corresponding to the final filament used (see Bonin et al., 2014).

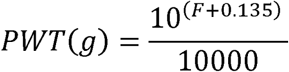

### Affective behavioral battery

Seventy-two hours after the final DiD session mice underwent three affective assays over three consecutive days in the fixed order: Elevated Zero Maze (day 1), Social Interaction (day 2), and Forced Swim Test (day 3).

### Elevated Zero Maze (EZM)

The EZM consisted of a circular track (outer diameter 61 cm; width 5 cm) elevated 51 cm above the floor and divided into four equal quadrants; two opposing quadrants were enclosed by walls (height 15 cm) and two were open. Each mouse was placed on the maze and allowed to explore for 5 minutes; behavior was video-recorded from above and hand-scored offline. Total time spent with the full body in open quadrants was summed as Total Open Arm Time and used as an index of anxiety-like behavior (Shepherd et al., 1994).

### Social Interaction (SI)

The SI apparatus comprised two connected Plexiglass chambers (30 x 30 x 30 cm) with a 2.5 cm central passage; a pencil cup was placed in each chamber. Three zones were defined per chamber: cup zone, inner zone (area from cup edge to half the box length), and outer zone. After a 5-minute habituation with no stimuli, the test mouse was temporarily removed while a naïve, younger, sex-matched conspecific was placed under one cup and a small tinfoil “object” under the other; positions were counterbalanced between trials. The test mouse was returned and allowed to explore for 8 minutes; video tracking (AnyMaze) quantified time spent in the inner zone around the naïve mouse as Social Interaction Time.

### Forced Swim Test (FST)

Four 4L beakers filled to ∼75% with water were placed in a row with dividers in between. The FST setup and procedure was largely based off previously published literature that utilize FST as a model of despair. Each mouse was placed in a beaker for a 6-minute trial; behavior was recorded. Latency to the first immobility bout (defined as ≥5 seconds of no active escape-directed movement) was scored. Stereotypic movement of a single paw used to maintain balance was included as immobility.

### Statistics

All statistics were calculated using Graph Pad Prism 10.4.2. Data was reported as mean +/- SEM. Data was collapsed by sex when analyzing between drug access or no drug access and separated by sex when analyzing within those that had drug access. Comparisons were analyzed using a 2-way ANOVA or mixed effects analysis, depending on the variables. Sidak’s multiple comparisons test was utilized post-hoc if there was a significant interaction between variables. Significance was defined as p ≤ 0.05. Individual z-scores for each behavior were calculated using 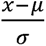 with (x) representing the individual value, (µ) the cohort mean and (σ) the cohort standard deviation. Average z-scores were calculated by taking the mean of an individual’s z-scores for escape jumps, wet-dog shakes and paw tremor for average withdrawal, and EZM, SI, and FST for average affective behavior. Values were sign-adjusted so positive values reflect exacerbated affective behavior (i.e., low open time, low social interaction, and low latency to immobility were transformed into high z-scores). For the FST, outliers were excluded using the ROUT method (Q = 1%). For cross-cohort exploratory analyses, behavioral measures were centered to each cohort’s water group, and difference-coded regression models and ordinal/hurdle models were implemented in Python; code is available upon request.

## Results

### Voluntary oxycodone two-bottle choice DiD does not elicit strong withdrawal

To establish whether mice would voluntarily consume dilute opioids without taste adulterants, we tracked total fluid intake, drug intake, and preference across four cohorts employing different opioids, session durations, concentration schedules, and assessed naloxone-precipitated (2 mg/kg, i.p.) somatic withdrawal on day 24.

In cohort 1, mice given access to oxycodone showed total fluid intake comparable to water controls across the 24-day paradigm, without any effect of oxycodone (Figure 2B), indicating that drug access did not suppress total fluid drinking. Oxycodone intake itself increased modestly as the concentration was increased from 0.1 to 1 mg/mL (main effect of day, p = 0.0063; Figure 2C), while preference for the oxycodone bottle remained stable between 40-60% throughout the paradigm in both sexes (Figure 2D). This voluntary intake, however, did not elicit physical dependence: escape jumps and paw tremors did not differ between oxycodone and water groups (Figures 2E-G). Oxycodone intake in this model was thus insufficient to produce robust naloxone-precipitated withdrawal.

**Figure 2.**
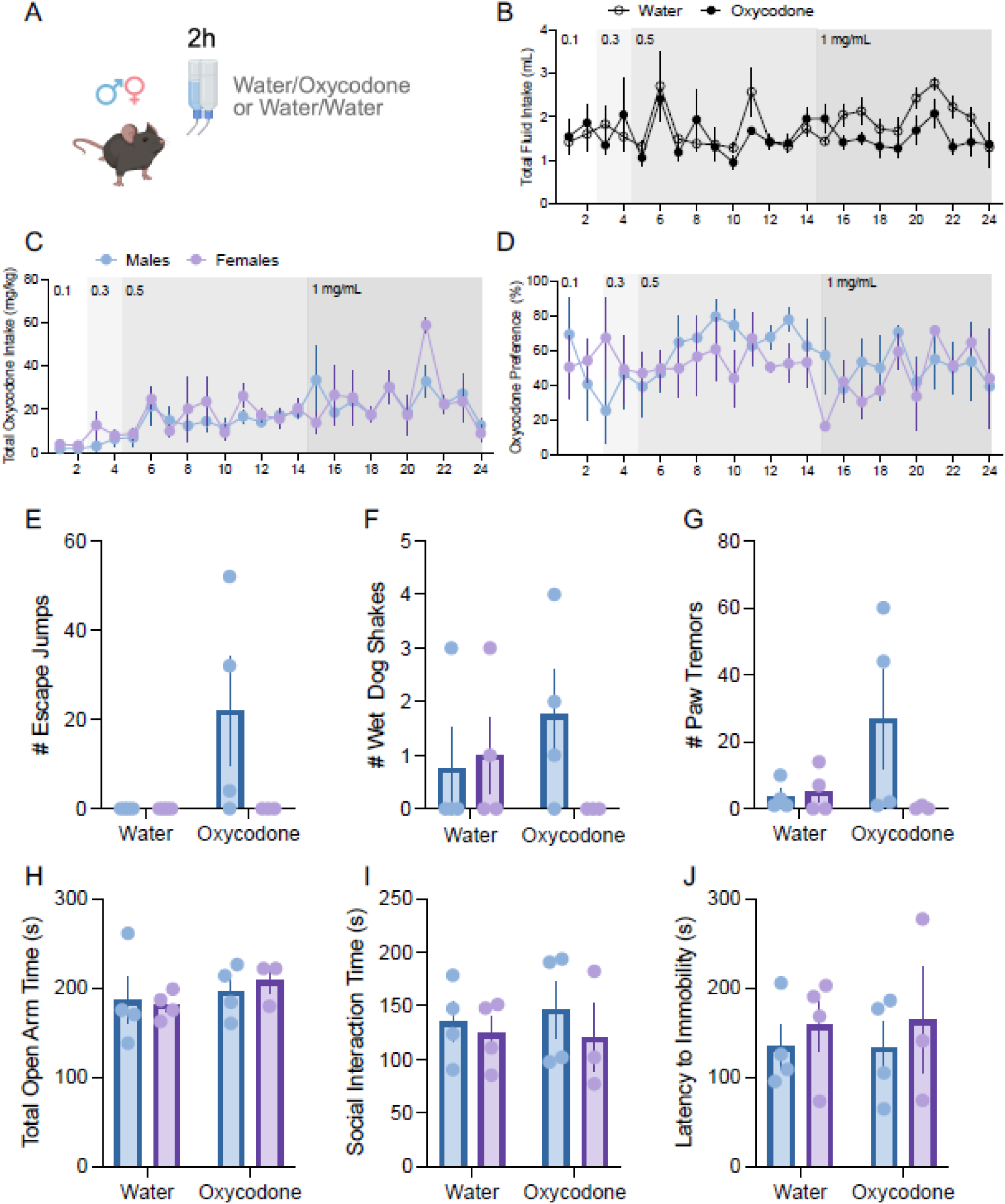
Longitudinal voluntary oxycodone drinking, naloxone precipitated withdrawal, and affective behavior. **A**. Schematic of the 2 hour/day, 24 day drinking in the dark (DiD) two bottle choice design; mice had daily access to either water/oxycodone (one bottle dilute oxycodone, one bottle water) or water/water (both bottles water). **B** Total daily fluid intake (mL; sum of both bottles) across the 24 day DiD paradigm for water (open circles) and oxycodone (filled circles) groups; mixed effects two way ANOVA revealed a significant effect of day; F(23,295) = 2.582, p < 0.0001. **C** Total oxycodone intake (mg/kg) in oxycodone access mice, separated by sex (males, blue; females, purple); mixed effects two way ANOVA showed a significant effect of day with Geisser–Greenhouse correction; F(3.403,17.90) = 5.446; ε = 0.1479. **D** Oxycodone preference (%) calculated as (oxycodone volume / total fluid volume) × 100 in oxycodone access mice, by sex; mixed effects two way ANOVA, ns. **E-G** Withdrawal signs scored after naloxone precipitated withdrawal; **E** number of escape jumps; **F** number of wet-dog shakes; **G** number of paw tremors; all three somatic measures analyzed by two way ANOVA, ns. **H** Total open arm time (s) in the Elevated Zero Maze; two way ANOVA, ns. **I** Total social interaction time (s) in the Social Interaction assay; two way ANOVA, ns. **J** Latency to immobility (s) in the Forced Swim Test; two way ANOVA, ns. Data are shown as mean ± SEM with individual datapoints overlaid. Sample sizes: water n = 8; oxycodone n = 8; males total = 8 (4/group); females total = 8 (4/group).

To evaluate whether chronic opioid exposure produced longer-lasting affective disturbances, mice were tested 72 hours after the final drinking session in the Elevated Zero Maze (EZM), Social Interaction (SI), and Forced Swim Test (FST). Composite affective z-scores were also calculated by averaging z-scores across the three assays for each animal. In Cohort 1 (oxycodone), no differences were detected between oxycodone and water groups in EZM open-arm time, SI time, or FST latency to immobility (Figures 2H-J).

### Short-access fentanyl DiD leads to withdrawal behaviors in male mice

In cohort 2, we provided mice with fentanyl access for 2 hours per day. Total fluid intake increased across day regardless of fentanyl access (p < 0.0001; Figure 3B). Fentanyl intake increased across the three concentrations steps (10, 20, 30 µg/mL; main effect of the day, p = 0.0364; Figure 3C), and fentanyl preference remained relatively constant across concentrations in both males and females (Figure 3D), indicating stable voluntary consumption at each dose. Withdrawal under this schedule occurred in males, as evidenced by increased wet-dog shakes (main effect of sex, p = 0.0131; Figure 3F). Escape jumps and paw tremors showed statistically significant sex difference (Figures 3E and 3G). There were no effects of fentanyl access or sex on EZM open arm time (3H) or immobility in the forced swim test (Figure 3J); however, female mice that had access to fentanyl displayed greater sociability than water controls. Overall, these data suggest that oral fentanyl can facilitate dependence and withdrawal, even under short access conditions.

**Figure 3.**
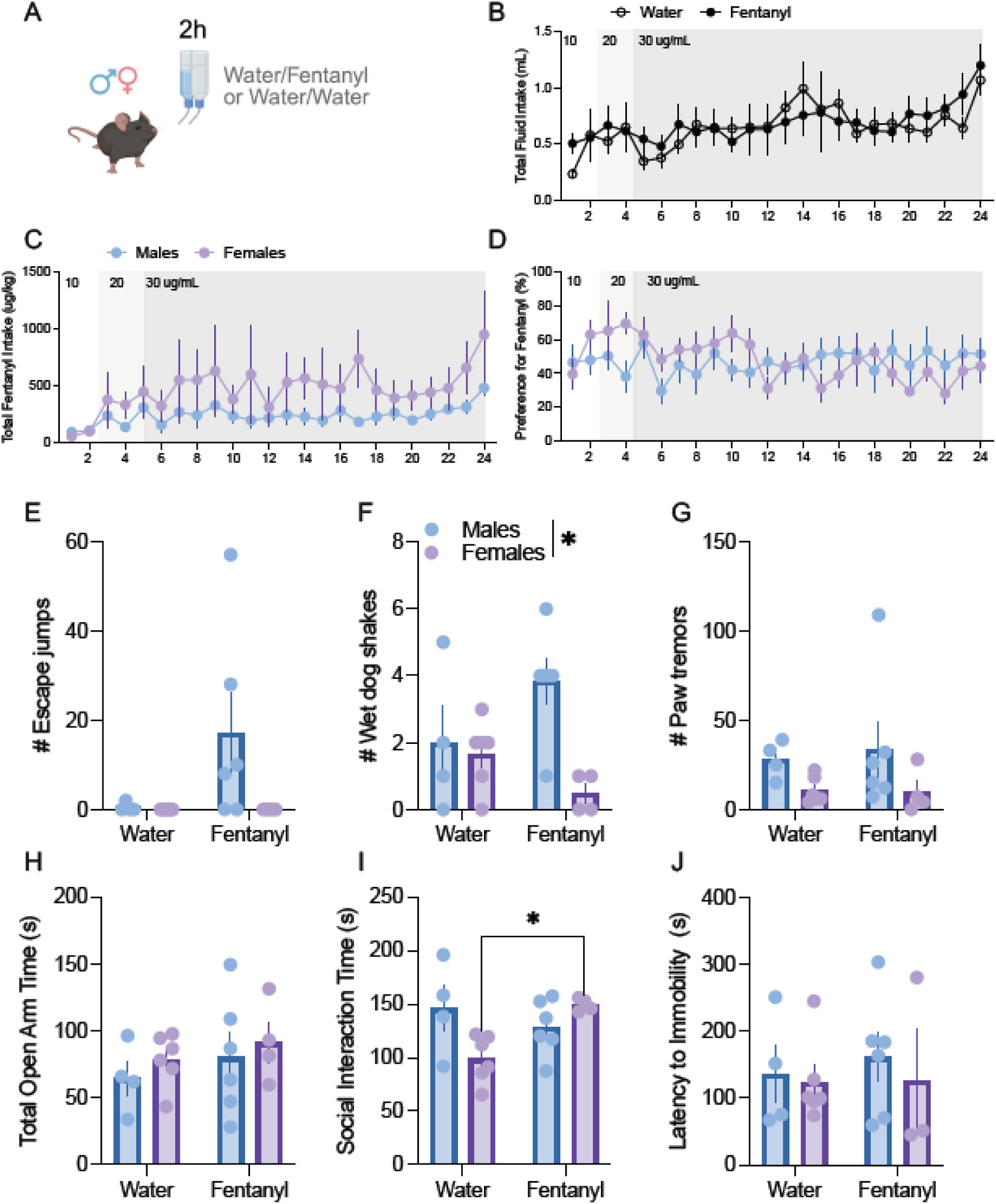
Longitudinal voluntary fentanyl drinking (2-hours access), naloxone precipitated withdrawal, and affective behavior. **A.** Schematic of the 2 hour/day, 24 day drinking in the dark (DiD) two bottle choice design; mice had daily access to either water/fentanyl (one bottle dilute fentanyl, one bottle water) or water/water (both bottles water). **B.** Total daily fluid intake (mL; sum of both bottles) across the 24 day DiD paradigm for water (open circles) and fentanyl (filled circles) groups; total fluid intake increased by day with no interaction with fentanyl; F(23,406) = 4.729, p < 0.0001 (**). **C.** Total fentanyl intake (µg/kg) in fentanyl-access mice, separated by sex (males, blue; females, purple); main effect of day with increased consumption over the paradigm; F(2.376, 18.91) = 3.749, p < 0.0. **D.** Fentanyl preference (%) calculated as (fentanyl volume / total fluid volume) × 100 in fentanyl-access mice, by sex; preference remained stable across the experiment and no aversion was observed in either sex. **E-G.** Withdrawal signs scored after naloxone-precipitated withdrawal; **E.** number of escape jumps; **F.** number of wet-dog shakes; **G.** number of paw tremors; analyses revealed a significant main effect of sex and a sex × fentanyl interaction for somatic signs overall; main effect of sex: F(1,16) = 7.783, p < 0.05; sex x fentanyl interaction: F(1,16) = 5.210, p < 0.05. **H.** Total open-arm time (s) in the Elevated Zero Maze. **I.** Total social interaction time (s) in the Social Interaction assay; sex x fentanyl interaction observed: F(1,16) = 7.603, p < 0.05 (*); post-hoc Sidak’s multiple comparisons test indicated females with fentanyl access spent more time in social interaction (Sidak, p < 0.05). **J.** Latency to immobility (s) in the Forced Swim Test. Data are shown as mean ± SEM with individual datapoints overlaid. Sample sizes: water n = 10; fentanyl n = 10; males: water 4, fentanyl 6; females: water 6, fentanyl 4 (for Forced Swim Test, females fentanyl n = 3).

### Longer-access fentanyl DiD leads to withdrawal behaviors in both sexes of mice

In Cohort 3, we maintained the concentration schedule for fentanyl throughout and extended access to 4h/day. Total fluid intake increased over days irrespective of fentanyl access (p = 0.0035; Figure 4B). As in Cohort 2, fentanyl intake increased across the concentration steps (main effect of day, p = 0.0084; Figure 4C), and preference again remained stable across concentrations in both sexes (Figure 4D). However, unlike Cohort 2, longer-access fentanyl DiD led to withdrawal in both males and, reflected in increases in both escape jumps (p = 0.0337; Figure 4E) and paw tremors (p = 0.0002; Figure 4G). Prolonged daily access from 2h to 4h at the same maximal concentration was therefore sufficient to strengthen withdrawal across both sexes. There was no effect of fentanyl DiD on EZM open arm time (Figure 4H), social interaction (4I), or FST immobility (Figure 4J), although there was an effect of sex on social interaction in this cohort.

**Figure 4.**
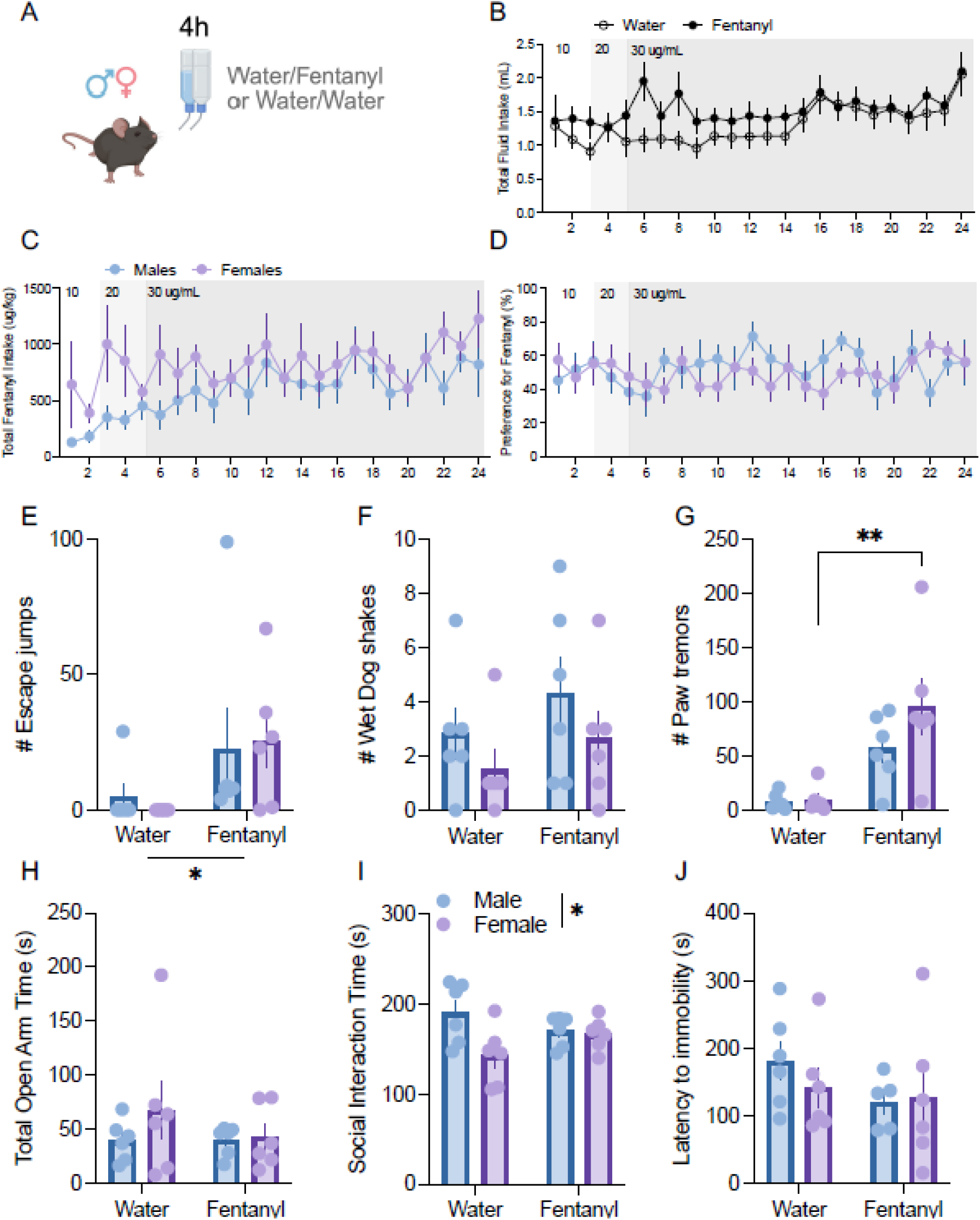
Longitudinal voluntary fentanyl drinking (4-hours access), naloxone precipitated withdrawal, and affective behavior (Cohort 3). **A**. Schematic of the 4 hour/day, 24 day drinking in the dark (DiD) two bottle choice design; mice had daily access to either water/fentanyl (one bottle dilute fentanyl, one bottle water) or water/water (both bottles water). **B**. Total daily fluid intake (mL; sum of both bottles) across the 24 day DiD paradigm for water (open circles) and fentanyl (filled circles) groups; main effect of day indicating increased drinking across the experiment; F(4.018,84.56) = 4.232, p < 0.01 (**). **C**. Total fentanyl intake (µg/kg) in fentanyl access mice, separated by sex (males, blue; females, purple); significant day × sex interaction; F(23,222) = 1.683, p < 0.05 (*). **D.** Fentanyl preference (%) calculated as (fentanyl volume / total fluid volume) × 100 in fentanyl access mice, by sex; preference remained stable and no aversion was observed in either sex. **E-G**. Withdrawal signs scored after naloxone precipitated withdrawal; **E** number of escape jumps; **F**. number of wet-dog shakes; **G.** number of paw tremors; fentanyl access increased escape jumps with a main effect of access F(1,20) = 5.202, p < 0.05 (*); fentanyl access increased paw tremors F(1,20) = 20.55, p < 0.001 (***); wet-dog shakes, ns. **H.** Total open arm time (s) in the Elevated Zero Maze; ns. **I**. Total social interaction time (s) in the Social Interaction assay; main effect of sex with females spending less time social interacting than males F(1,20) = 5.496, p = 0.05 (*). **J**. Latency to immobility (s) in the Forced Swim Test; ns. Data are shown as mean ± SEM with individual datapoints overlaid. Sample sizes: water n = 12; fentanyl n = 12; males: water 6, fentanyl 6; females; water 6, fentanyl 6 (for Forced Swim Test, fentanyl males n = 5).

### Higher concentrations of fentanyl lead to greater withdrawal in female mice

The most pronounced and sex-divergent phenotype emerged in Cohort 4, where mice underwent a high-concentration fentanyl escalation (10-100 µg/mL) over 4h/day. In this cohort, fentanyl decreased total fluid intake. Although total fluid intake still rose across days (p < 0.0001), fentanyl-access mice drank less overall than water controls (main effect of fentanyl, p = 0.0109; Figure 5B). Despite this, fentanyl intake climbed sharply across the escalation (main effect of day, p < 0.0001), with females consuming more than males (main effect of sex, p = 0.0206; day x sex interaction, p = 0.0074; Figure 5C). Females also sustained high fentanyl preference throughout, while males’ preference declined once concentrations exceeded 60 µg/mL (Figure 5D). In the withdrawal test, fentanyl-exposed females showed greater escape jumps (Figure 5E) and paw tremors (Figure 5G). There was also an interaction between sex and fentanyl that affected wet-dog shakes (Figure 5F). Despite the increased fentanyl intake and withdrawal in females, we observed minimal evidence for long-term affective disturbances (Figure 5H-J), as in previous cohorts.

**Figure 5.**
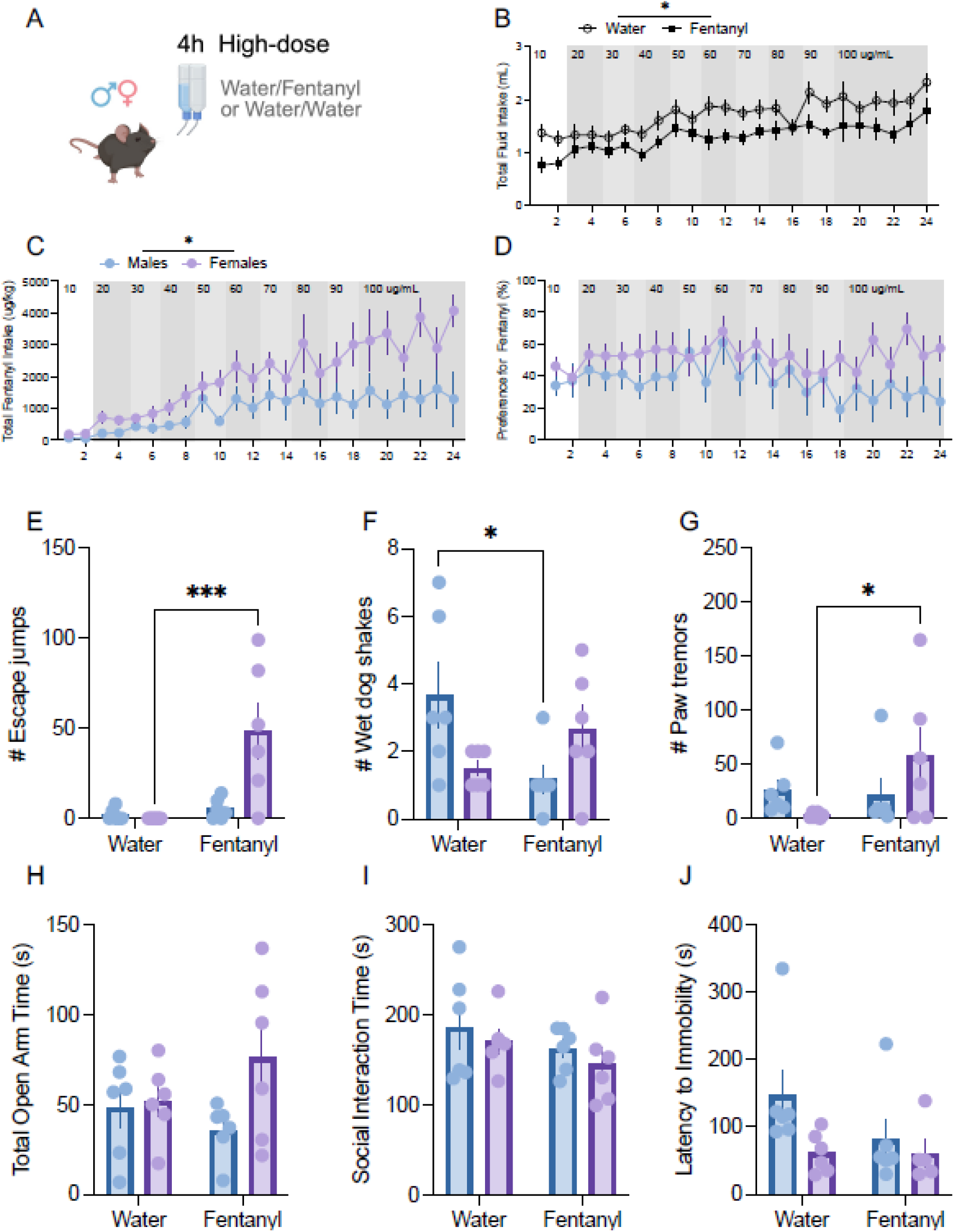
Longitudinal voluntary fentanyl drinking (4-hours access, high-dose), naloxone precipitated withdrawal, and affective behavior. **A.** Schematic of the 4 h/day high-concentration two-bottle choice drinking in the dark (DiD) paradigm across 24 days. Mice received daily access to either water/fentanyl (one bottle fentanyl solution, one bottle water) or water/water (both bottles water). **B.** Total daily fluid intake (mL; combined intake from both bottles) across the 24 day DiD paradigm for water (open circles) and fentanyl (filled circles) groups. Fluid intake increased across days (F_(23, 496)_ = 8.567, ****p < 0.0001), and fentanyl-access mice consumed less total fluid overall compared to controls (F_(1, 22_) = 7.742), *p < 0.05). **C.** Total fentanyl intake (µg/kg) across the 24 day DiD paradigm in fentanyl-access mice separated by sex (males, blue; females, purple). Total fentanyl intake increased significantly as an effect of day (F_(4.328, 42.52)_ = 9.176, ****p < 0.0001) and females exhibited greater fentanyl intake across the drinking period compared to males (F_(1, 10)_ = 7.538, *p < 0.05). There was a significant interaction between day x sex (F_(23, 226)_ = 1.950, **p < 0.01). **D.** Preference for fentanyl (%) calculated as (fentanyl volume / total fluid volume) × 100 in fentanyl-access mice by sex. Females displayed greater fentanyl preference relative to males across the drinking period. **E–G.** Somatic withdrawal signs assessed following precipitated withdrawal. **E.** Number of escape jumps. **F.** Number of wet-dog shakes. **G.** Number of paw tremors. Fentanyl access increased escape jumps and paw tremors relative to water controls, while wet-dog shakes were not significantly altered. The animals with access to fentanyl experienced more escape jumps (F_(1, 20)_ = 11.29, **p < 0.01) and females overall experienced more escape jumps than males (F_(1, 20)_ = 7.120, *p < 0.05). There was an interaction of fentanyl x sex (F_(1, 20)_ = 8.571, **p < 0.01) and female animals with access to one bottle of fentanyl demonstrated significantly more escape jumps (Sidak’s multiple comparisons test, ***p < 0.001). Furthermore, there was a significant interaction demonstrated between fentanyl x sex for wet-dog shakes (F_(1, 20)_ = 8.231, **p < 0.01) and male animals with access to one bottle of fentanyl demonstrated significantly less wet-dog shakes (Sidak’s multiple comparisons test, *p < 0.05). **H.** Total open arm time (s) in the Elevated Zero Maze; no significant effect of fentanyl access was observed. **I.** Total social interaction time (s) in the Social Interaction assay; no significant group differences were observed. **J.** Latency to immobility (s) in the Forced Swim Test; no significant effect of fentanyl access was detected. Data are presented as mean ± SEM with individual datapoints overlaid. Asterisks indicate significant effects (*p < 0.05, ***p < 0.001). Sample sizes: water n = 12; fentanyl n = 12; males: water 6, fentanyl 6; females: water 6, fentanyl 6. (for Forced Swim Test, fentanyl females n = 5).

### Minimal persistent affective behavioral alterations following chronic opioid intake

To facilitate comparisons across groups and assess potential individual differences that may present across behavioral endpoints, we generated composite z-scores for withdrawal and for affective behaviors. This analysis generally supported the previous conclusions. In Cohort 1, the composite affective z-scores showed no effect of oxycodone access (Figure 6C). In cohort 3 (Figure 6H), we observed a main effect of fentanyl on the withdrawal Z-score, suggesting that fentanyl access led to withdrawal in both male and female mice. In Cohorts 2 and 4 we observed sex by fentanyl interactions driven by increased withdrawal in male and female mice, respectively. Furthermore, there was no effect of fentanyl or sex by fentanyl interactions that mediate affective behavior during abstinence in this model. Furthermore, we observed no effects of fentanyl access on mechanical sensitivity in the Von Frey test (Supplemental Figure 1). Together, these results indicate that despite producing strong somatic withdrawal, voluntary fentanyl intake in this paradigm was not consistently associated with persistent affective-like disturbances 72 hours after the final drinking session.

**Figure 6.**
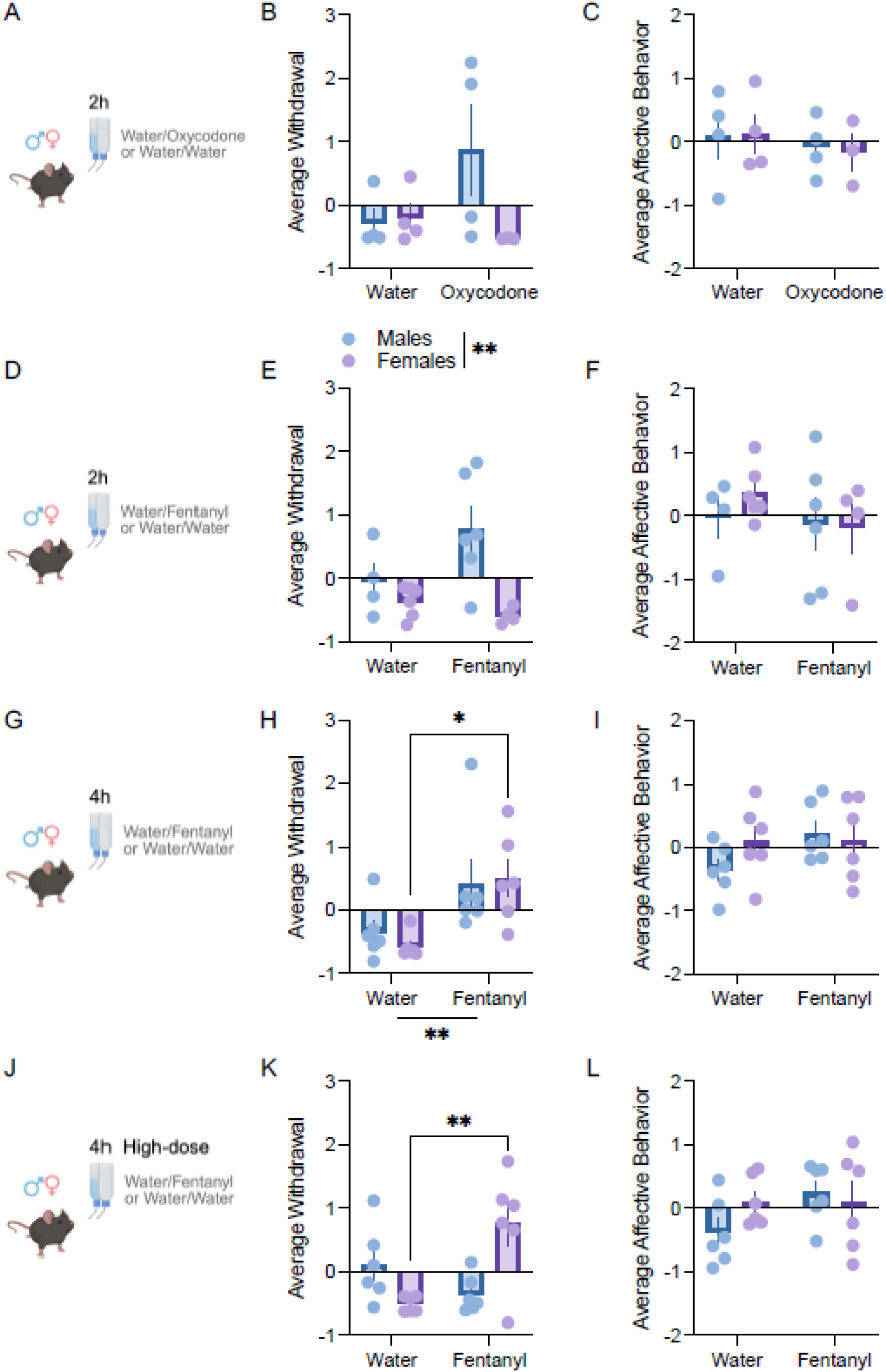
Average z scored withdrawal and affective behavior across oxycodone and fentanyl drinking cohorts. AC. Cohort 1:2 h oxycodone drinking paradigm. **A.** Schematic of the 2 h/day two-bottle choice drinking in the dark (DiD) paradigm in which mice received access to either water/oxycodone or water/water. **B.** Average z-scored withdrawal behavior in males (blue) and females (purple), including individual animal z-scores for escape jumps, paw tremors, and wet-dog shakes. Two male oxycodone-drinking animals exhibited elevated withdrawal scores relative to the remainder of the oxycodone group, although this effect was not consistent across the male oxycodone-drinking cohort. **C.** Average z-scored affective behavior, including individual animal z-scores for total open arm time in the Elevated Zero Maze (EZM), latency to immobility in the Forced Swim Test (FST), and total social interaction (SI) time. No significant group differences were observed. **D-F. Cohort 2: 2 h fentanyl drinking paradigm. D.** Schematic of the 2 h/day fentanyl two-bottle choice DiD paradigm. **E.** Average z-scored withdrawal behavior among males and females. Male mice displayed significantly greater withdrawal behavior relative to females; main effect of sex, F(1,16) = 10.97, p < 0.01 (**). **F.** Average z-scored affective behavior including EZM, FST, and SI measures. No significant effects were detected. **G-I. Cohort 3: 4 h fentanyl drinking paradigm. G.** Schematic of the 4 h/day fentanyl two-bottle choice DiD paradigm. **H.** Average z-scored withdrawal behavior among males and females. Mice with fentanyl access exhibited significantly greater withdrawal behavior compared to water controls; main effect of fentanyl access, F(1,20) = 12.55, p < 0.001 (**). **I.** Average z-scored affective behavior including EZM, FST, and SI measures. No significant effects were observed. **J-L. Cohort 4: 4 h high-concentration fentanyl drinking paradigm. J.** Schematic of the 4 h/day high-concentration fentanyl two-bottle choice DiD paradigm. **K.** Average z-scored withdrawal behavior among males and females. There was a significant fentanyl access × sex interaction, F(1,20) = 15.16, p < 0.001 (*). Females with fentanyl access displayed significantly greater withdrawal behavior relative to water controls (Sidak’s multiple comparisons test, p < 0.01. **L.** Average z-scored affective behavior including EZM, FST, and SI measures. No significant differences were detected. Data are presented as mean ± SEM with individual datapoints overlaid. Withdrawal composite z-scores were calculated from escape jumps, paw tremors, and wet-dog shakes. Affective composite z-scores were calculated from Elevated Zero Maze, Forced Swim Test, and Social Interaction behavioral measures.

### Mechanical nociceptive thresholds were unaltered following opioid exposure

In all cohorts, mechanical sensitivity was assessed immediately after naloxone-precipitated withdrawal using the von Frey assay to determine paw withdrawal thresholds (PWT). Within each cohort, standard group comparisons revealed no significant effects of oxycodone or fentanyl access, or sex, on PWT (Supplementary Figure 1A-D), indicating that the DiD regimens did not produce clear mechanical hyperalgesia at the time of testing.

To compare across fentanyl cohorts while accounting for baseline differences, we used an ordered logit model with difference-coded cohorts and centering to each cohort’s water group. This analysis detected an effect of cohort on PWT (global χ² test p ≈ 0.008), driven primarily by a small shift between Cohort 2 and Cohort 3, but found no reliable effect of fentanyl treatment or sex. In a separate ordinal model limited to fentanyl-access mice only, higher log-transformed fentanyl consumption showed a weak, non-significant trend toward lower PWT (p ≈ 0.07). Overall, these findings suggest that trans-cohort differences in PWT largely reflect baseline protocol differences rather than consistent fentanyl-induced mechanical hyperalgesia, and that any consumption-related changes in mechanical sensitivity are subtle at best.

### Multidimensional behavioral profiles visualized by radar/spyder plots

To provide an integrated view of behavioral outcomes across withdrawal, affective, and nociceptive domains, we generated radar plots depicting mean z-scores for each measure within experimental groups (Supplementary Figure 2). Cohort 1 (oxycodone) exhibited modest, inconsistent elevations in withdrawal-associated z-scores with minimal changes in affective or nociceptive measures (Supplementary Figure 2A). Cohort 2 (2 h fentanyl) displayed elevated withdrawal-related z-scores, particularly in males, with limited affective alterations (Supplementary Figure 2B). Cohort 3 (4 h fentanyl) showed broader increases in withdrawal-related z-scores across both sexes while affective and mechanical sensitivity measures remained relatively stable (Supplementary Figure 2C). Cohort 4 (high-concentration fentanyl) produced the most pronounced behavioral phenotype, characterized by elevated withdrawal-associated z-scores, especially in females, with comparatively smaller effects on affective behavior and nociception (Supplementary Figure 2D).

To probe these patterns further, we modeled withdrawal, affective, and nociceptive measures across cohorts using difference-coded regression models centered to each cohort’s water group. For paw tremors, fentanyl increased tremors when moving from the short (2 h) to the long (4 h) access protocol (Cohort 2 vs 3), followed by a relative decrease when moving from the standard-dose to the high-concentration 4 h protocol (Cohort 3 vs 4); a small sex by treatment interaction was also detected, though effect sizes and confidence intervals suggested this was weak and may reflect noise. Because many animals exhibited zero escape jumps, we additionally examined individual fentanyl consumption in relation to jumping behavior among fentanyl-access mice: higher log-transformed fentanyl consumption was associated with a greater number of jumps, particularly in females (sex x consumption interaction, p = 0.03), suggesting that the magnitude of voluntary intake more closely tracks withdrawal severity in females than in males. By contrast, no comparable dose-response relationships were detected for affective measures (EZM, SI, FST) or for PWT, consistent with the absence of robust fentanyl- or consumption-related effects on these domains.

Together, these multidimensional profiles and trans-cohort analyses reinforce that session duration and dose escalation modulate the magnitude and sex-specificity of opioid withdrawal phenotypes, with voluntary fentanyl intake more closely tracking withdrawal severity in females, whereas affective-like and nociceptive endpoints remain comparatively unaffected by opioid exposure.

## Discussion

### Voluntary fentanyl self-administration can elicit precipitated withdrawal

Across four cohorts, male and female C57BL/6J mice readily consumed dilute oxycodone and fentanyl in a two-bottle DiD paradigm without taste adulterants. In three of the four cohorts, drug availability did not reduce total fluid consumption relative to water-only controls, and drug preference remained stable at ∼40-60% across multiple oxycodone and fentanyl concentrations. These findings indicate that mice will voluntarily consume translationally relevant opioids in a simple, low-cost oral paradigm that does not require surgical catheterization or sweeteners, extending previous work on oral oxycodone and fentanyl models (Downs et al., 2024; Mavrikaki et al., 2017; Peretz-Rivlin et al., 2025; Zhang et al., 2014).

### Experimental parameters dictate bidirectional sex differences in withdrawal

With oxycodone, voluntary intake led to minimal physical dependence since only a few oxycodone drinking males displayed escape jumps, and there was no group level effect on composite withdrawal scores. This is consistent with the relatively modest dose range achieved here, and suggest that higher concentrations of oxycodone or longer session durations regimens may be required to produce robust precipitated withdrawal in a two-bottle choice paradigm. Fentanyl, by contrast, produced clear naloxone-precipitated withdrawal, and withdrawal severity depended on both session duration and concentration. When fentanyl access was limited to 2 h/day with a maximal concentration of 30 µg/mL (Cohort 2), withdrawal was present but predominantly expressed in males, who showed more somatic signs than females, despite comparable intake between sexes. Increasing session duration to 4 h/day (Cohort 3) eliminated the sex difference such that both males and females displayed elevated escape jumps, paw tremors, and composite z-scores. Thus, for the same fentanyl concentration ramp, extending access from 2h to 4h transformed a male-biased withdrawal profile into a robust, sex-independent fentanyl withdrawal phenotype.

When fentanyl dose was further increase to 100 µg/mL under 4 h/day access (Cohort 4), we observed withdrawal behaviors only in female mice. Fentanyl-access animals consumed less total fluid than water controls, and within the fentanyl group, males showed lower total fentanyl intake and a gradual decline in fentanyl preference once concentrations exceeded ∼60 µg/mL. In contrast, females maintained stable preference and continued to increase total fentanyl intake as a function of rising concentration. This divergence in intake was mirrored in withdrawal since females but not males displayed increased escape jumps, paw tremors, and composite z-scores. Dose-response analyses restricted to fentanyl-access animals indicated that higher total fentanyl consumption modestly increased the probability of exhibiting escape jumps and, among animals that jumped, was associated with greater jump counts, particularly in females. These data are consistent with emerging reports that females can show higher oral opioid intake than males under certain conditions (Downs et al., 2024; Zanni et al., 2021) and suggest that at higher concentrations males begin to exhibit avoidance, whereas females tolerate or overcome the pharmacological effects and/or bitter taste to maintain intake. These observations suggest that as cumulative fentanyl exposure increases, females are more likely to reach a threshold sufficient for prominent physical dependence, whereas males may limit intake or shift to avoidance at higher concentrations.

### Limited persistent affective alterations despite clear somatic withdrawal

We next asked whether chronic oral oxycodone or fentanyl exposure produced longer-lasting changes in behaviors related to anxiety, depression, or sociability. Across all cohorts, composite affective z-scores derived from EZM, SI, and FST did not differ between drug and water groups. These results suggest that, under the present exposure conditions and time frame, voluntary oral oxycodone and fentanyl induce somatic withdrawal without consistent changes to affective-like behaviors. This contrasts with some drug injection and vapor fentanyl studies in which extended access has yielded anxiety- and depression-like phenotypes (Moussawi et al., 2020; Marchette et al., 2023), and with intravenous fentanyl self-administration models in which anxiety-like behavior emerge only after a period of abstinence (Chen et al., 2025). It is possible that affective aftereffects would emerge under longer withdrawal intervals, more severe dependence, or in the context of additional stressors or learning tasks, such as fear extinction, as reported in other oral fentanyl paradigms (Downs et al., 2024).

### Mechanical nociceptive sensitivity is largely preserved across fentanyl regimens

Mechanical nociception, assessed by von Frey thresholds immediately after withdrawal, was not robustly altered by oxycodone or fentanyl access. Ordinal regression models across fentanyl cohorts detected an effect of cohort on PWT that appeared to reflect baseline differences between cohorts rather than systematic fentanyl effects, and ordinal models restricted to fentanyl-access mice revealed at most a weak, non-significant trend toward lower thresholds with higher fentanyl consumption. Thus, within the dose and timing parameters used here, voluntary oral fentanyl did not produce clear mechanical hyperalgesia. The absence of mechanical sensitivity is noteworthy given the literature describing hyperalgesia following repeated opioid exposure, particularly following high parenteral dosing or sustained infusions (Delgado et al., 2023; Koponen and Forget, 2022). In addition, hyperalgesia in fentanyl-dependent mice has been shown to emerge during spontaneous withdrawal windows of 36-48 hours post-cessation (Chen et al., 2025). By contrast, we assessed mechanical hypersensitivity at a much earlier timepoint. We believe this variable, along with the moderate cumulative exposure and voluntary nature of intake in our oral paradigm, likely contributed to the absence of robust mechanical hyperalgesia in the present study. Future work could systematically explore these parameters, along with different sensory modalities.

### Translational relevance, advantages, and limitations of the model

The present work advances the toolbox for modeling opioid use and dependence by demonstrating that a simple, two-bottle DiD protocol can support voluntary oral oxycodone and fentanyl intake. The paradigm can be tuned by adjusting access duration and dose escalation to generate measurable, physical withdrawal, with or without sex differences in intake. Compared with intravenous or vapor self-administration, the oral DiD paradigm reduces surgical burden, training time, and equipment cost, is easily scalable to larger cohorts, and better approximates oral misuse patterns seen with prescription opioids. The two-bottle configuration additionally disentangles fluid homeostasis from drug-directed preference, allowing assessment of motivational allocation toward opioid intake.

Despite the strengths of this approach, several limitations should be considered. First, the oxycodone regimen used here produced minimal withdrawal, so the present study primarily informs fentanyl dependence. Stronger oxycodone exposure (e.g, higher concentrations, longer access, or extended duration) will be required to determine whether analogous parameters can lead to oxycodone dependence in this model. Second, while we observed clear sex differences in intake and withdrawal at high fentanyl doses, the underlying mechanisms were not examined and remain to be elucidated. Third, the behavioral battery was limited to three assays at a single time point three days after the last drinking session; more extensive affective testing or longer withdrawal intervals might reveal additional consequences of chronic oral oxycodone and fentanyl. Finally, although we used centering and difference-coded regression to account for cohort baselines and protocol changes, the three fentanyl cohorts differed in both session duration and dose range, which constrains strict causal attribution.

## Conclusions and future directions

In summary, this study establishes a voluntary two-bottle Drinking-in-the-Dark model of oral oxycodone and fentanyl self-administration in male and female mice and defines exposure parameters under which fentanyl, but not oxycodone, produces robust precipitated withdrawal. Longer access sessions and higher fentanyl concentrations markedly increase withdrawal severity, and high-dose escalation preferentially unmasks stronger dependence in females. In contrast, affective-like behaviors and mechanical nociceptive thresholds remain largely preserved under the conditions tested and are only weakly related, if at all, to individual fentanyl consumption. These features make the present paradigm a practical and translationally relevant platform for studying the circuit and molecular mechanisms of voluntary opioid dependence and for testing candidate interventions. Future work can leverage this model to examine how specific brain regions and cell types encode oral fentanyl intake and withdrawal, to probe protracted withdrawal and relapse-like behavior, and to investigate sex-specific vulnerability and resilience factors in opioid use and dependence.

## Data availability

Raw data will be made available on an online repository prior to publication.

## Acknowledgements

We thank members of the Joffe lab for helpful discussions and feedback.

## Author Contributions

MCA: writing – original draft, data curation, visualization; SFB: investigation, writing – original draft; NDJ: investigation; LHN: formal analysis; MEJ: conceptualization, writing-review and editing, supervision, and funding acquisition.

---

**Supplementary Figure 1.**
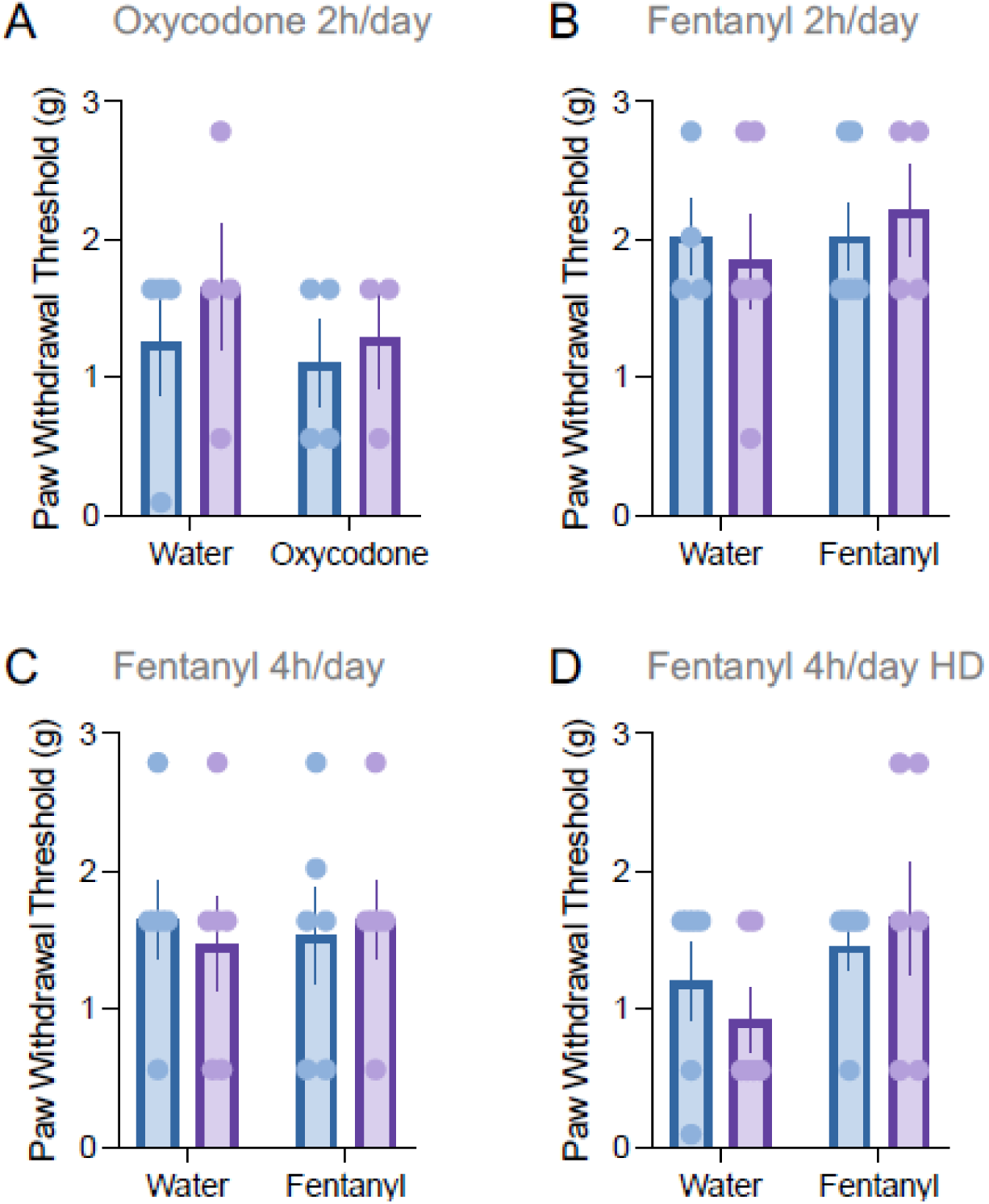
Mechanical nociceptive thresholds following oxycodone or fentanyl drinking across cohorts. A. Cohort 1: Oxycodone 2 h/day. Paw withdrawal threshold (g) measured using the von Frey assay in water- and oxycodone-access mice separated by sex (males, blue; females, purple). No significant effects of oxycodone access or sex were observed. **B. Cohort 2: Fentanyl 2 h/day.** Paw withdrawal threshold (g) measured using the von Frey assay in water- and fentanyl-access mice separated by sex. No significant differences in mechanical sensitivity were detected between groups. **C. Cohort 3: Fentanyl 4 h/day.** Paw withdrawal threshold (g) measured using the von Frey assay following 4 h/day fentanyl drinking. No significant effects of fentanyl access or sex were observed. **D. Cohort 4: Fentanyl 4 h/day high-concentration (HD).** Paw withdrawal threshold (g) measured using the von Frey assay in mice exposed to the high-concentration fentanyl drinking paradigm. Mechanical sensitivity did not significantly differ between water- and fentanyl-access groups. Data are presented as mean ± SEM with individual datapoints overlaid. Von Frey testing was used to assess mechanical nociceptive sensitivity following completion of the drinking paradigms.

**Supplementary Figure 2.**
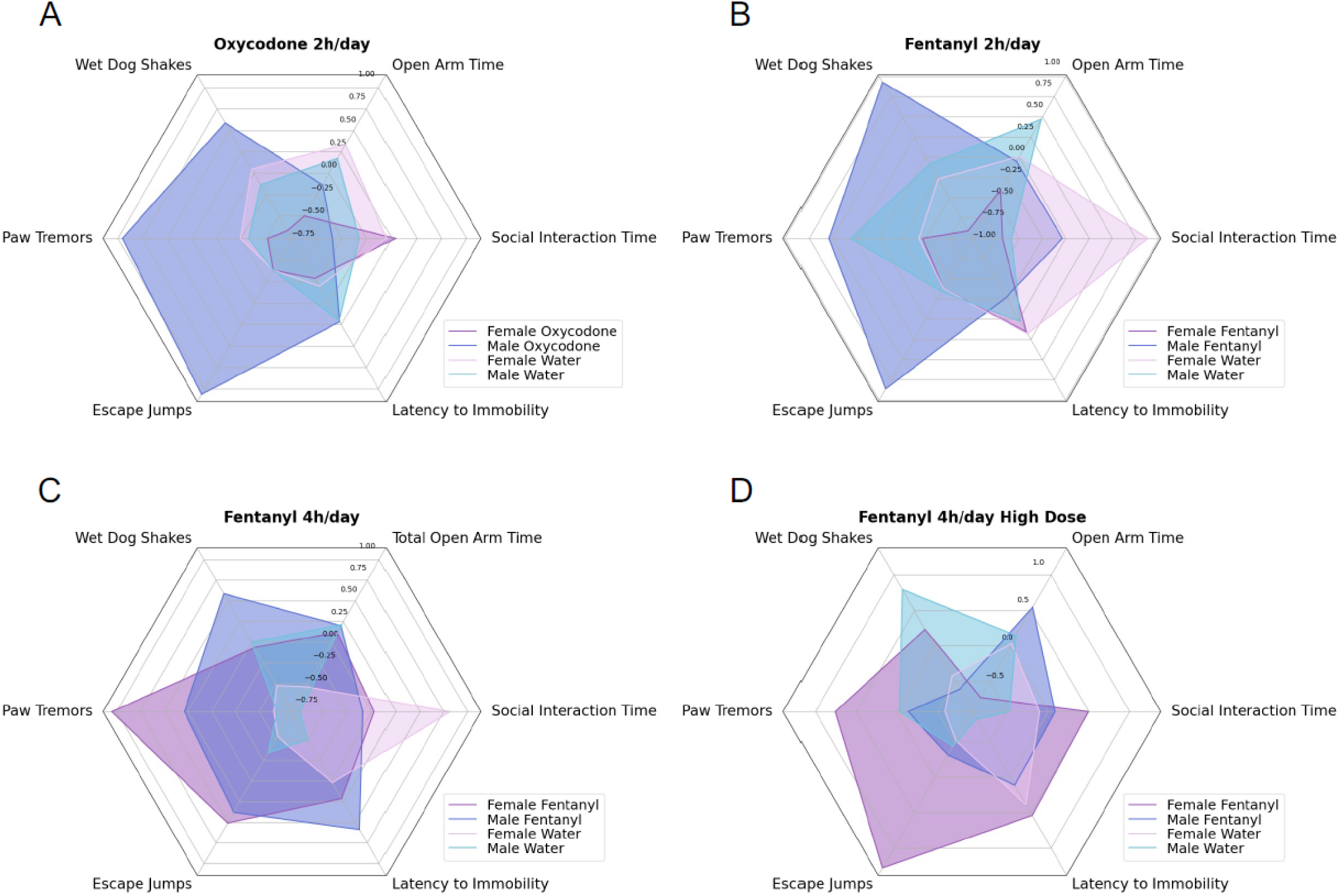
Spider plot visualization of behavioral z-scores across oxycodone and fentanyl drinking cohorts. Spider plots depict the mean z-score for each behavioral measure across experimental groups within each cohort following completion of the drinking paradigms. Individual behavioral measures included somatic withdrawal signs (escape jumps, paw tremors, wet-dog shakes), affective-like behaviors (Elevated Zero Maze open arm time, Forced Swim Test latency to immobility, and Social Interaction time), and mechanical sensitivity measured using the von Frey assay. Z-scores were calculated for each behavioral measure and averaged within experimental groups to allow comparison of multidimensional behavioral profiles across cohorts. **A. Cohort 1: Oxycodone 2 h/day.** Oxycodone-access mice exhibited modest increases in withdrawal-associated behavioral z-scores relative to water controls, while affective-like and nociceptive measures remained largely comparable between groups. **B. Cohort 2: Fentanyl 2 h/day.** Fentanyl-access mice demonstrated elevated z-scores for withdrawal-associated behaviors compared to controls, with limited alterations in affective or nociceptive measures. **C. Cohort 3: Fentanyl 4 h/day.** Mice with extended fentanyl access displayed broader increases in withdrawal-related behavioral z-scores relative to water controls, whereas affective-like and mechanical sensitivity measures remained relatively stable. **D. Cohort 4: Fentanyl 4 h/day high-concentration.** High-concentration fentanyl exposure produced the most pronounced behavioral phenotype, characterized by elevated withdrawal-associated z-scores, particularly in females, with comparatively smaller effects on affective-like behavior and nociceptive sensitivity. Spider plots are intended to provide a multidimensional visualization of group-level behavioral profiles across cohorts and assays rather than individual statistical comparisons.

**Supplementary Figure 3.**
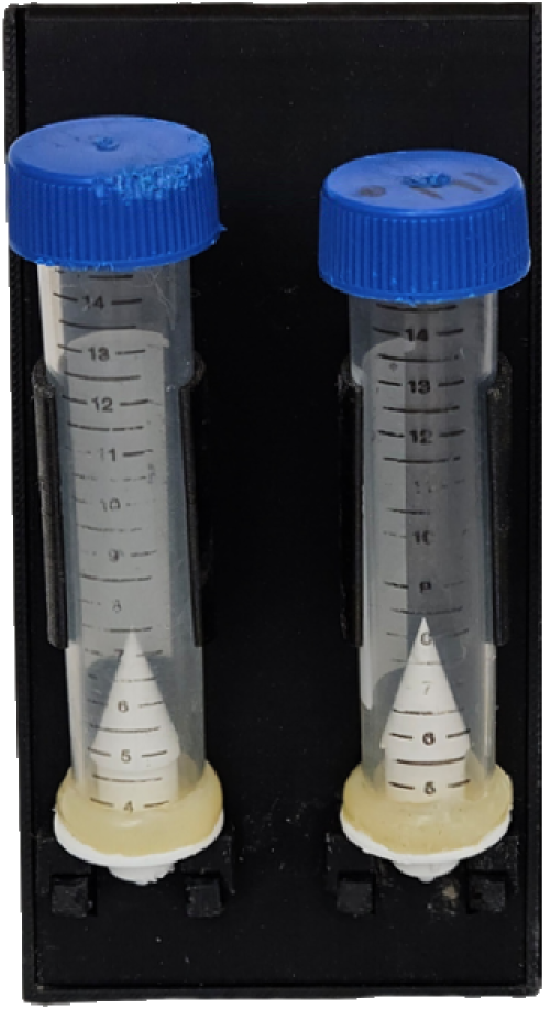
Representative image of the 2BC sipper setup.

